# The adaptive stochasticity hypothesis: modelling equifinality, multifinality and adaptation to adversity

**DOI:** 10.1101/2023.05.02.539045

**Authors:** Sofia Carozza, Danyal Akarca, Duncan Astle

**Author notes:** **Corresponding authors:** Sofia Carozza, Danyal Akarca. Co-lead authors (Sofia Carozza, Danyal Akarca).

## Abstract

Neural phenotypes are the result of probabilistic developmental processes. This means that stochasticity is an intrinsic aspect of the brain as it self-organizes over a protracted period. In other words, while both genomic and environmental factors shape the developing nervous system, another significant—though often neglected—contributor is the randomness introduced by probability distributions. Using generative modelling of brain networks, we provide a framework for probing the contribution of stochasticity to neurodevelopmental diversity. To mimic the prenatal scaffold of brain structure set by activity-independent mechanisms, we start our simulations from the medio-posterior neonatal rich-club (Developing Human Connectome Project; *dHCP, n* = 630). From this initial starting point, models implementing Hebbian-like wiring processes generate variable yet consistently plausible brain network topologies. By analyzing repeated runs of the generative process (> 10^7^ simulations), we identify critical determinants and effects of stochasticity. Namely, we find that stochastic variation has a greater impact on brain organization when networks develop under weaker constraints. This heightened stochasticity makes brain networks more robust to random and targeted attacks, but more often results in non-normative phenotypic outcomes. To test our framework empirically, we evaluated whether stochasticity varies according to the experience of early-life deprivation using a cohort of neurodiverse children (Centre for Attention, Learning and Memory; *CALM n* = 357). We show that low socioeconomic status predicts more stochastic brain wiring. We conclude that stochasticity may be an unappreciated contributor to relevant developmental outcomes, and make specific predictions for future research.

## INTRODUCTION

Human brain structure is the result of a complex and dynamic interplay among various constraints. Foremost among them are the genomic information children receive from their parents and the environment in which they grow up^1^. In the literature, these two factors—together with the interaction between them—are often credited with explaining the whole of phenotypic variation in brain development across the population^2^. However, this overlooks a critical fact: that development unfolds stochastically^3,4^.

Stochasticity refers to the fact that biological development is probabilistic, rather than deterministic; there is an element of intrinsic randomness or noise in the relationship between earlier and later states^5^. This appears to be an integral—rather than artefactual—feature of many developmental processes. At the cellular level, due to non-linear interactions between molecules and entropy-induced variation, identical biochemical processes result in different outcomes across individuals^6^. In neurodevelopment itself, stochasticity is particularly operative through randomness in the transcription and translation of key proteins^7^, axonal outgrowth,^8,9^ and dynamics of spontaneous neuronal firing and synaptic transmission^10,11^. These processes converge to produce stochastic influences on the morphology of macroscopic brain regions. The relative importance of stochasticity as a contributor to phenotypic outcomes likely varies by domain. Some features—such as the volume of the brain stem—are highly heritable and appear to be tightly governed by genetic constraints^12^, while others—such as the microstructure of association tracts—are only weakly heritable and vary greatly even between genetically identical individuals^13^. While some of this non-heritable variation is due to differential environmental exposures, which engage experience-dependent neural processes^14,15^, other variation is likely attributable to inherent stochasticity within brain development.

As developmental stochasticity heightens intra-individual variability, its contribution to phenotypic outcomes is an adaptive feature that favors species success in environmental challenges^6^. Evolutionary pressures may therefore have favored a *heightened* role for stochastic developmental processes in harsh and uncertain environments. Exposure to unpredictability early in life is a robust predictor of later behavior across species^16^, including cognitive and emotional outcomes in humans^17– 21^. This is thought to reflect adaptive responses to ancestral cues or to statistical learning of environmental changes^22^. Could ontogenetic stochasticity account for some of this pathway? In other words, could stochastic processes in neural development mediate adaptation to unpredictable early-life environments?

Despite being an inherent feature of neural development, across multiple levels of analysis, stochasticity is largely neglected in empirical studies. One reason for this lacuna could be the difficulty of successfully separating stochastic effects from unknown deterministic effects^3^, let alone manipulating it to evaluate the magnitude of its contribution to a particular outcome.

A promising path toward addressing this gap lies in computational modelling, which permits a quasi-experimental approach to understanding the emergence of neural phenotypes. One such method, generative network modelling, probabilistically simulates realistic whole-brain networks^23,24^. In this model, nodes within a developing network form connections based on an economic trade-off between two parameters: the *cost* of forming a connection versus the *value* that connection may bring. Crucially, this trade-off can be constrained to varying extents by modulating parameter magnitude. Highly constrained simulations minimize stochasticity within the generative process, which may lead to limited phenotypic variability. In contrast, weaker wiring constraints allow for greater randomness from one step of the generative process to the next, which may thereby produce greater phenotypic variation. Thus, in this model of the brain, stochasticity is an integral and manipulatable element of development.

In this work, we explored the contributions of stochasticity to whole-brain organization by analyzing repeated runs of the generative network model^24–26^. Starting from a prenatal scaffold of brain architecture obtained from the Developing Human Connectome Project (*dHCP, n* = 630) we generated over ten million plausible brain networks. We analyzed the simulations to answer the following questions: Does developmental stochasticity lead to variability in brain network outcomes? Are some connectomes more sensitive to the effects of heightened developmental stochasticity than others? What advantage might stochastic development confer on brain networks? Through this work, we produce a framework for understanding determinants of variability in brain network organization.

To test the empirical plausibility of our theoretical framework, we then simulated the connectomes of a sample of neurodiverse children (Centre for Attention, Learning and Memory; *CALM, n* = 357). Children who grew up in early socioeconomic deprivation showed macroscopic brain networks that appear to organize more stochastically. We propose this finding may reflect an adaptive connectomic response to unpredictable features of the early environment. We conclude with specific predictions and recommendations for future investigation that follow from our framework.

## RESULTS

To simulate the formation of brain network connectivity, we employed a generative network model. This model is increasingly used in computational neuroscience to simulate highly plausible brain networks^25–33^. The model does this by adding connections iteratively based on dynamic economic negotiations between their costs and topological values^23,24^:

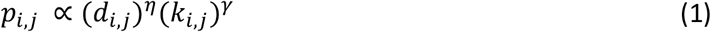

The *d*_*i,j*_ term represents the cost of a connection between two given nodal regions *i* and *j*, which we approximated using the Euclidean distance between the regions. The *k*_*i,j*_ term represents the topological value of this connection, which we estimated using the normalized overlap in connectivity between regions *i* and *j. p*_*i,j*_ refers to the overall the probability of forming a fixed binary connection between nodes *i* and *j*, and is proportional to the parametrized multiplication of costs and values. Two wiring parameters, *η* and *γ*, respectively scale the contributions of each term to wiring probability. In other words, by varying *η* and *γ*, it is possible to impose different constraints on the developing network.

The generative model begins from an initial minimal scaffold of connectivity. To root our simulations in a realistic representation of neonatal brain structure, we reconstructed a core rich club network from data collected from the Developing Human Connectome Project (*dHCP*; see **Methods** and **Supplementary Figures 1-2**). Because connections form within this initial scaffold probabilistically, the generative model is intrinsically stochastic. This means there may not be a clear one-to-one relationship between wiring parameters and outcomes. For example, running the model multiple times using the same *η* and *γ* parameters may lead to highly dissimilar phenotypes, whilst running the model with different *η* and *γ* parameters may lead to highly similar phenotypes (**Figure 1a**).

**Figure 1.**
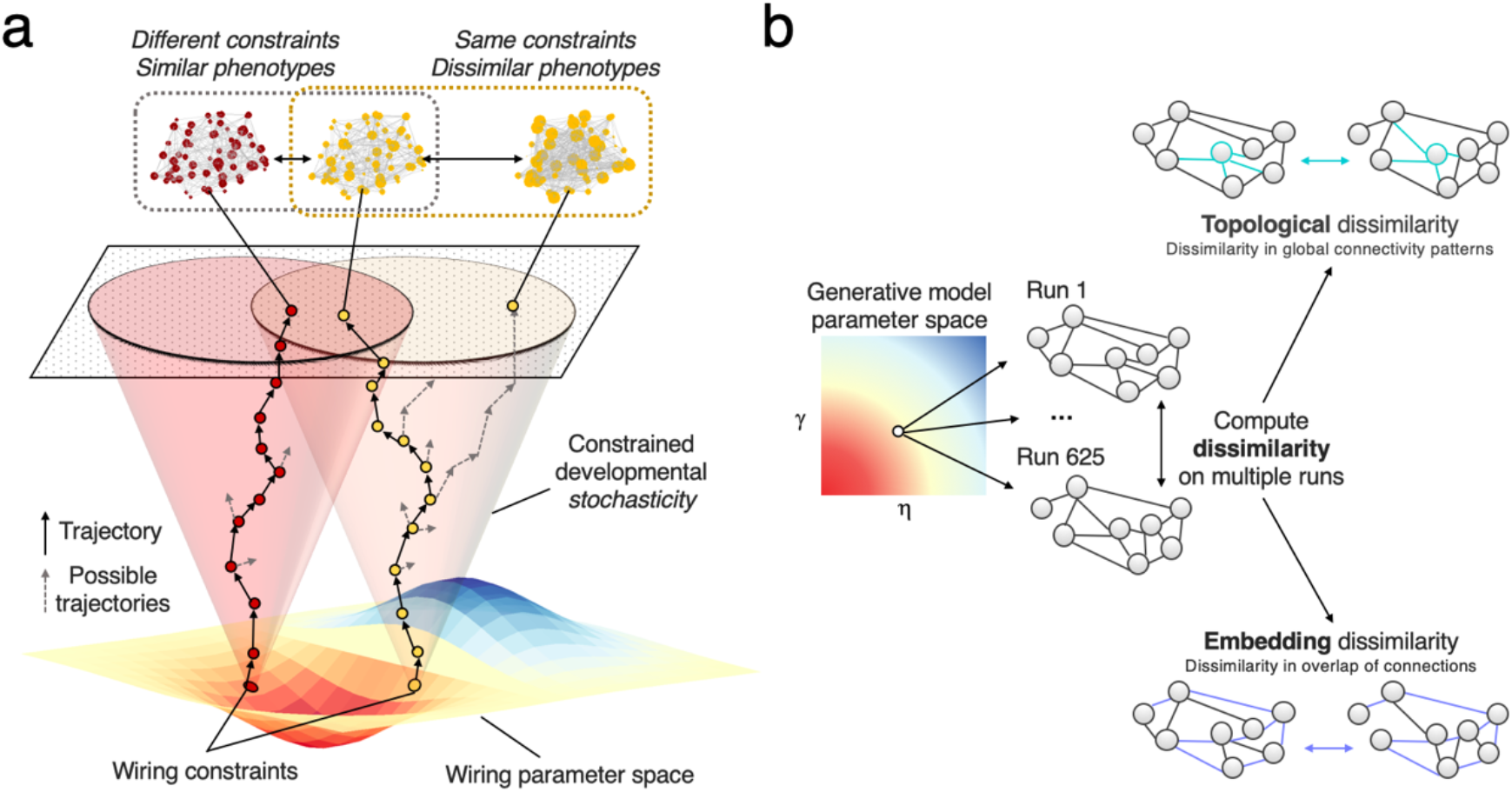
Schematic of dissimilarity procedure. (a) A schematic illustration of how stochasticity in the developing simulation leads to variable outcomes. For example, the same wiring constraints can lead to quite dissimilar outcomes (i.e., multifinality, top right) and different wiring constraints can lead to similar outcomes (i.e., equifinality, top left). (b) For each parameter combination, we ran 625 repeated simulations. To calculate the dissimilarity between these network outcomes, two measures were produced. The first was a topological dissimilarity measure, which calculated the dissimilarity in global network topology between each pairwise combination of network outcomes at each parameter combination (top). The second was an embedding dissimilarity measure, which calculated the dissimilarity in edge existence between each pairwise combination of network outcomes at each parameter combination (bottom).

### Experiment 1 – The effect of constraints on multifinality

We first set out to determine the relationship between constraints on development and variability in network outcomes. This is equivalent to *multifinality*, a known developmental principle by which the same developmental history can result in diverse phenotypes^34^ (**Figure 1a**, top right). Specifically, we asked: does changing the wiring constraints systematically manipulate its intrinsic stochasticity, therefore leading to altered variability in observed brain organization?

We addressed this question by undertaking repeated runs of the generative model with identical wiring parameters, at multiple combinations of *η* and *γ*. In order to quantify the resulting variability in network outcomes, we calculated two measures: topological and embedding dissimilarity (**Figure 1b**; see **Methods: Experiment 1**). Topological dissimilarity captures how different networks are to each other in terms of their organization; a low topological dissimilarity indicates similar topology across repeated runs of the simulation, whilst a high topological dissimilarity indicates multifinality. In contrast, embedding dissimilarity tests whether networks resemble one another in the precise location of their connections. A low embedding dissimilarity means that repeated runs of the simulation share many specific connections between the same nodes, whilst a high embedding dissimilarity indicates multifinality.

We first explored the topological dissimilarity across the wiring parameter space (**Figure 2a**). We found that constraints on wiring are in fact systematically related to multifinality in brain network organization. However, *η* and *γ* do not contribute to topological dissimilarity in the same way. Namely, *γ*, which guides the extent to which regions prefer connecting to regions with similar neighborhoods, has a much larger influence on network stochasticity (R^2^, 76.3%) compared to *η*, which penalizes long-distance connections (R^2^, 0.1%) (**Figure 2b**). In **Figure 2c**, this result is presented schematically, illustrating that softer *γ* constraints drive networks to be topologically highly dissimilar across runs, while fewer differences emerge when the *γ* parameter is larger in magnitude.

**Figure 2.**
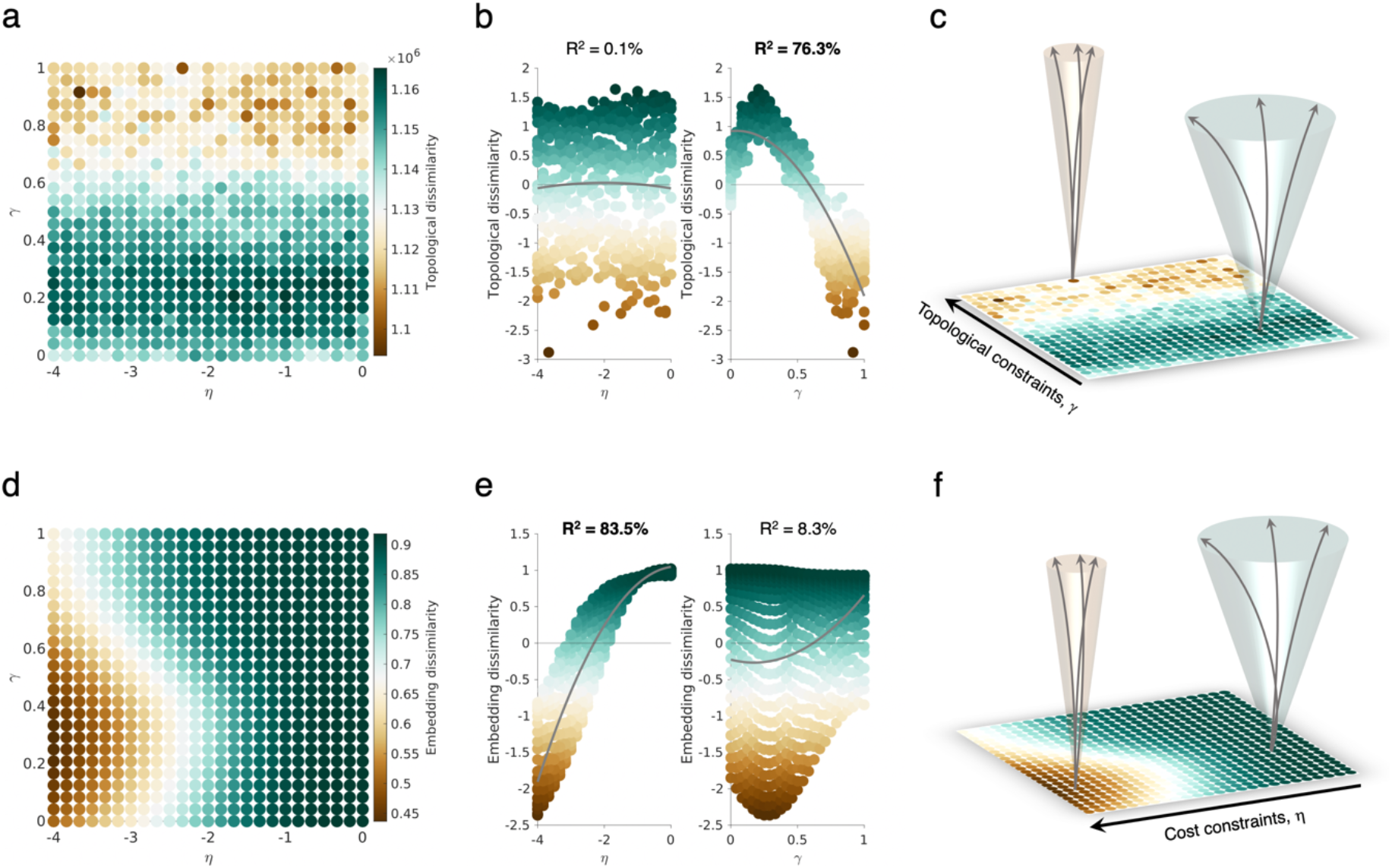
Weaker brain wiring constraints increase multifinality in network outcomes. (a) The topological dissimilarity landscape is given across the wiring parameter space. Green corresponds to highly topologically dissimilar networks while brown corresponds to low topological dissimilarity. (b) A scatter plot of the topological dissimilarity as a function of the wiring parameters shows that *γ* most drives topological dissimilarity. (c) Variable outcomes are more likely with weaker topological constraints on the network development (highlighted by the light green wider funnel) and vice versa for highly constrained networks (highlighted by the light brown narrower funnel). (d) The embedding dissimilarity landscape is given across the wiring parameter space. Green corresponds to highly dissimilar networks in terms of embeddings while brown corresponds to low embedding dissimilarity (e) A scatter plot of the topological dissimilarity as a function of the wiring parameters shows that *η* most drives embedding dissimilarity. (f) Variable outcomes are more likely with less embedding constraints on the network development (highlighted by the light green wider funnel) and vice versa for highly constrained networks (highlighted by the light brown narrower funnel).

We then explored the embedding dissimilarity across the wiring parameter space. As shown In **Figure 2d**, an opposite effect occurs, in which *η* has a much larger influence (R^2^, 83.5%) compared to *γ* (R^2^, 8.3%) (**Figure 2e**). The schematic in **Figure 2f** illustrates this, showing that softer *η* constraints drive networks to more dissimilar topological states, while fewer differences emerge when the *η* parameter is larger in (negative) magnitude.

These findings can be interpreted quite straightforwardly from the wiring equation. As *γ* determines the extent to which nodes with similar connectivity profiles wire with each other, a strong *γ* favours a similar topological arrangement across runs. Across repeated runs, the same organizational features will emerge. When *η* has a large magnitude, the simulation favors the formation of local short connections—these are more consistently present across multiple runs, increasing the embedding similarity. Thus, across multiple runs of the simulation, the same brain wiring parameters can produce networks that vary in organization (topological dissimilarity) and in the precise location of their connections (embedding dissimilarity), depending upon the severity of each of the two wiring constraints.

Overall, weaker wiring constraints results in more heterogeneity in brain network outcomes, while stronger wiring constraints reduce stochasticity and limit the range of possible outcomes.

### Experiment 2 – The effect of noise on multifinality

Our first experiment revealed that softening wiring constraints leads to greater multifinality in the organization and spatial localization of the connectome. But can variation in network outcomes be increased by enhancing the noisiness of network development itself? In other words, does upregulating developmental stochasticity increase multifinality? And if so, *when* does this intervention have the greatest influence on network outcomes?

To answer these questions, we heightened the stochasticity of the simulations at different stages of network development. To do this, we allowed the model to choose 5% of total connections completely at random either at the start of, halfway through, or in the final steps of the generative process (termed *early, middle*, and *late* respectively) (see **Methods: Experiment *2***).

We first examined the effect of heightened developmental noise on topological dissimilarity. Our results indicate that the *later* the noise is injected into the developmental simulation, the greater the increase in topological dissimilarity across multiple runs of the same parameters (**Figure 5a-c, Supplementary Figure 4**, ANOVA F_2,1872,_ = 134.44, *p* = 2.774 × 10^−55^; early M = 1.242 × 10^3^, SD = 6.685 × 10^3^; middle M = 8.212 × 10^3^, SD = 9.826 × 10^3^; late M = 8.296 × 10^3^, SD = 9.348 × 10^3^). The effect was predominantly present at higher values of *γ* (see **Supplementary Figure 4 i-k**), where the simulations are more strongly driven by the wiring rule. Thus, the topology of networks that are following a more deterministic, rule-based approach to development are more sensitive to the effects of injecting noise.

**Figure 3.**
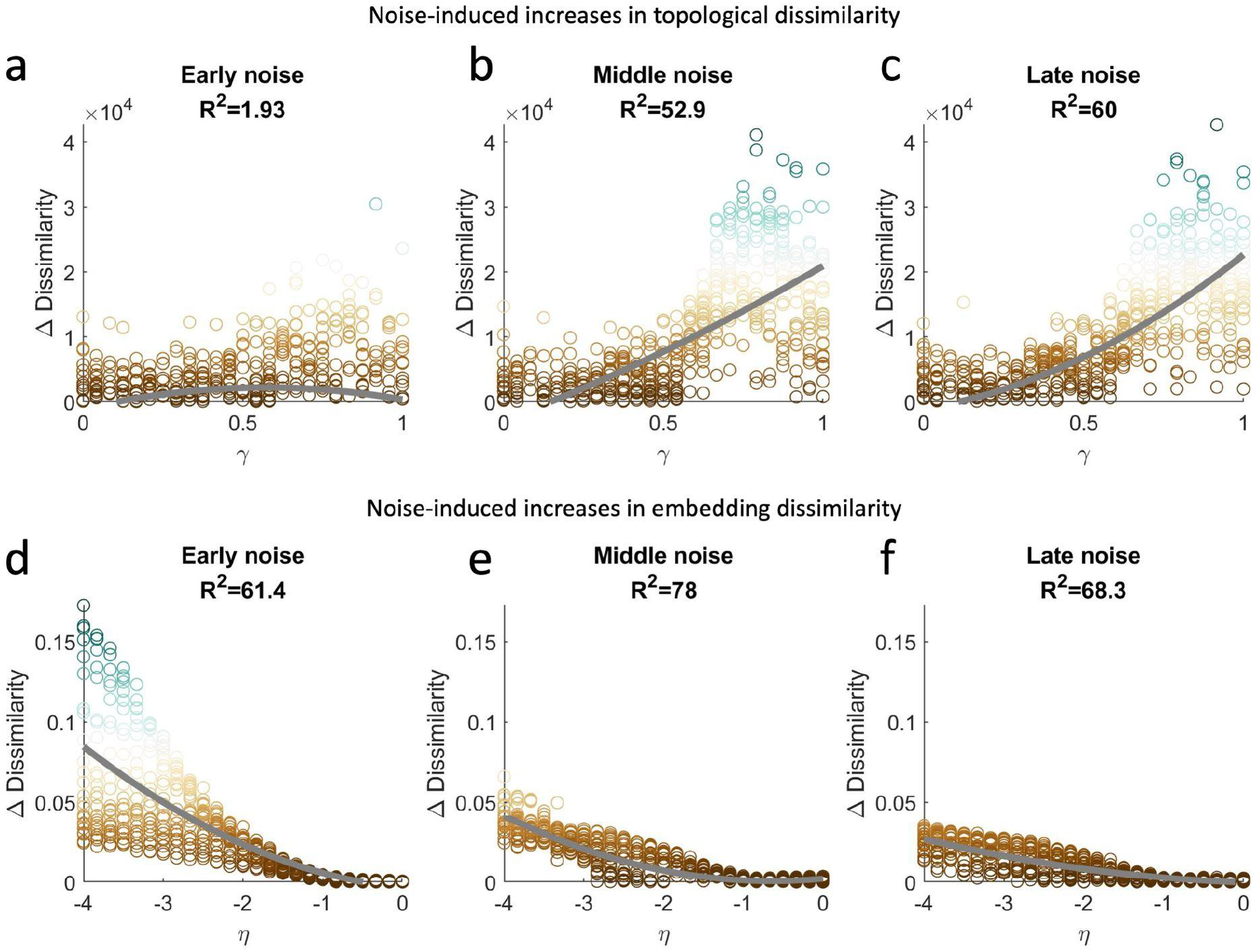
Upregulating developmental noise increases variability in network outcomes. Injecting stochasticity (a) early, (b) middle, or (c) late in the generative process increased the topological dissimilarity across repeated runs of each parameter combination, with middle and late noise exerting a greater impact (ANOVA F_2,1872,_ = 134.44, *p* = 2.774 × 10^−55^) especially at higher values of *γ*. Injecting stochasticity (d) early, (e) middle, or (f) late in the generative process also increased the embedding dissimilarity, with early noise exerting a greater impact (ANOVA F_2,1872,_ = 140.92, *p* = 9.792 × 10^−58^) especially at lower values of *η*.

**Figure 4.**
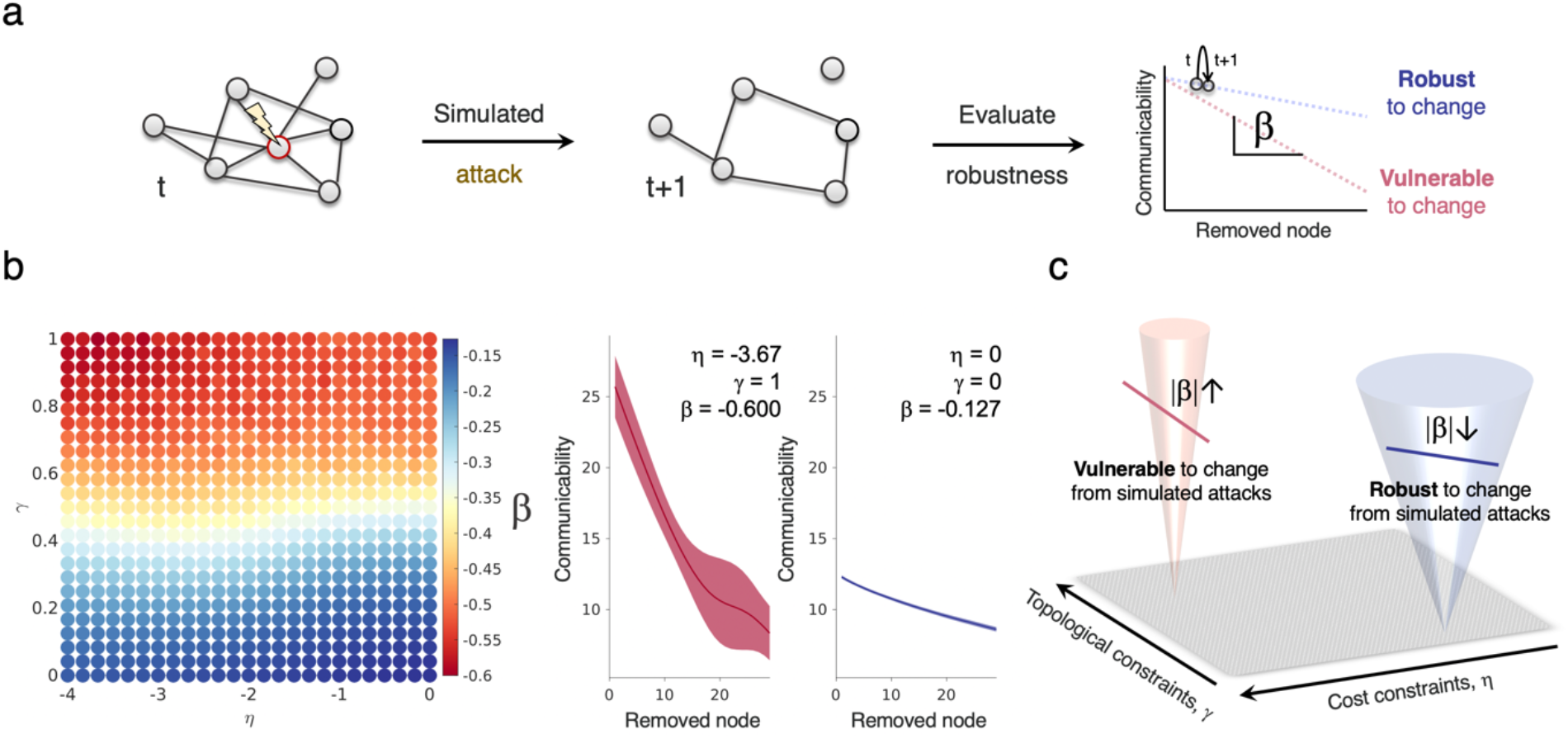
Weaker wiring constraints confer relative resilience to simulated attacks. (a) A schematic demonstration of the robustness testing protocol. Robustness is estimated by quantifying how much network communicability decreases upon the removal of network nodes. (b) The *β* coefficient computed from a targeted attack regime, which preferentially removes network hubs, across the parameter space. More constrained networks (top left of the landscape) are less robust, as the gradient of change is greater. Weakly constrained networks (bottom right of the landscape) exhibit less change. To the right of the landscape, the communicability trajectories of networks with the least (left) and most (right) robustness to change are presented. (c) A schematic showing that weakly constrained networks (which achieve more multifinality, as indicated by the funnel width), are relatively more robust to attack.

**Figure 5.**
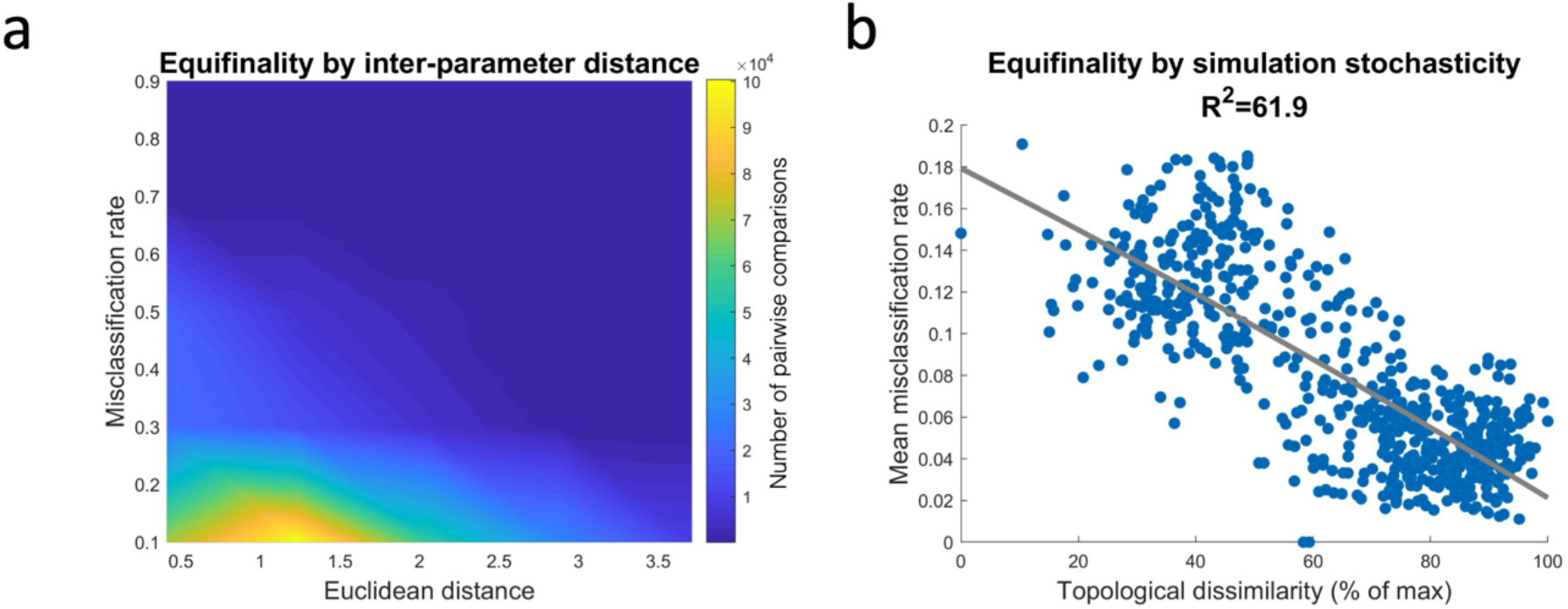
Similar wiring constraints and deterministic development lead to equifinality in network outcomes. A support vector machine (SVM) was trained to distinguish simulations run with different parameters. The SVM sought to correctly classify the runs of the simulations using their global statistics. The mean misclassification rate refers to the mean proportion of the sample that was incorrectly classified across all pairwise comparisons, and was obtained using cross validation. (a) A density plot of equifinality, measured using the misclassification rate of the SVM, by the Euclidean distance between the two parameter combinations. Each point represents a pair-wise comparison of the global statistics of two multi-run simulations. (b) The equifinality exhibited by a parameter combination, indexed using the mean misclassification rate of the SVM, by the topological dissimilarity of that simulation. The mean was taken over all pairwise simulation comparisons.

We next evaluated the impact on embedding dissimilarity. Interestingly, we find the opposite phenomenon, showing that the *earlier* noise is injected into the developmental simulation, the greater the increase in embedding dissimilarity across runs (**Figure 5d-f, Supplemental Figure 5**, ANOVA F_2,1872,_ = 140.92, *p* = 9.792 × 10^−58^; early M = 0.0298, SD = 0.0246; middle M = 0.0126, SD = 0.0144; late M = 0.0103, SD = 0.0101). Injecting noise results in a greater increase in outcome variability when models are further from the origin of the parameter space (**Supplemental Figure 5i-l**). This greater impact of early stochasticity on the final layout of connections is consistent with the importance of the initial scaffold upon which network development unfolds.

In summary, the timing of heightened ontogenetic stochasticity shapes its impact on resulting network outcomes. Weaker wiring constraints (i.e., low magnitude parameters) broadly protect against the impact of injecting noise at any time point during development. Networks that are developing under stronger wiring constraints are more sensitive, but this effect depends on the timing of the intervention. Namely, early stochasticity has little effect on topological dissimilarity but greatly increases embedding dissimilarity, whilst late stochasticity greatly increases topological dissimilarity with a negligible impact on embedding. This indicates that topological characteristics of the network can recover from temporary increases in developmental noise, given sufficient time, but that the precise identity and locations of the connections that form depend strongly on the initial scaffold.

### Experiment 3 – The effect of constraints on robustness

Our first two experiments revealed that networks with weaker wiring constraints exhibit greater multifinality but are less sensitive to the effects of temporary increases in developmental noise. In contrast, highly constrained networks with exhibit less multifinality but are more vulnerable to such interventions, depending on their timing.

As an evolved system, the brain is best understood in light of selective pressures that shaped its features across evolutionary history^35^. This includes the brain’s potential to reach multiple phenotypic outcomes. Why might this be advantageous? One possibility is that higher stochasticity within development, which leads to multifinality, confers robustness to external perturbation. This phenomenon has been observed in developmental systems in biology at multiple levels^36^.

To examine this possibility, we tested whether higher stochasticity within the simulation confers robustness to targeted attacks on nodes with high levels of connectivity. **Figure 4a** provides a schematic overview of the experimental procedure. In short, by measuring the resilience of each network’s communicability to the removal of nodes, we obtained a measure of relative robustness to external perturbation (see **Methods: Experiment 3**). The smaller the change in response to simulated attacks, the greater the robustness.

Our findings indicate that networks with higher constraints on their topology are more vulnerable to targeted attacks relative to networks with lesser constraints (**Figure 4b**). In **Supplementary Figure 6** we show this also true for a random attack regime. Given that strong wiring constraints produce topologically invariant networks, these may rely more strongly on core hub nodes, resulting in a proportionally larger drop in network communication when such nodes are removed. However, it is important to note that this large drop still leaves the networks at a higher *absolute* capacity for communication than networks that develop with weaker wiring constraints. Thus, networks developing more deterministically are more vulnerable to change in relative terms, while retaining a more advantageous topology than networks that develop more stochastically. For a schematic presentation of this finding, see **Figure 4c**.

**Figure 6.**
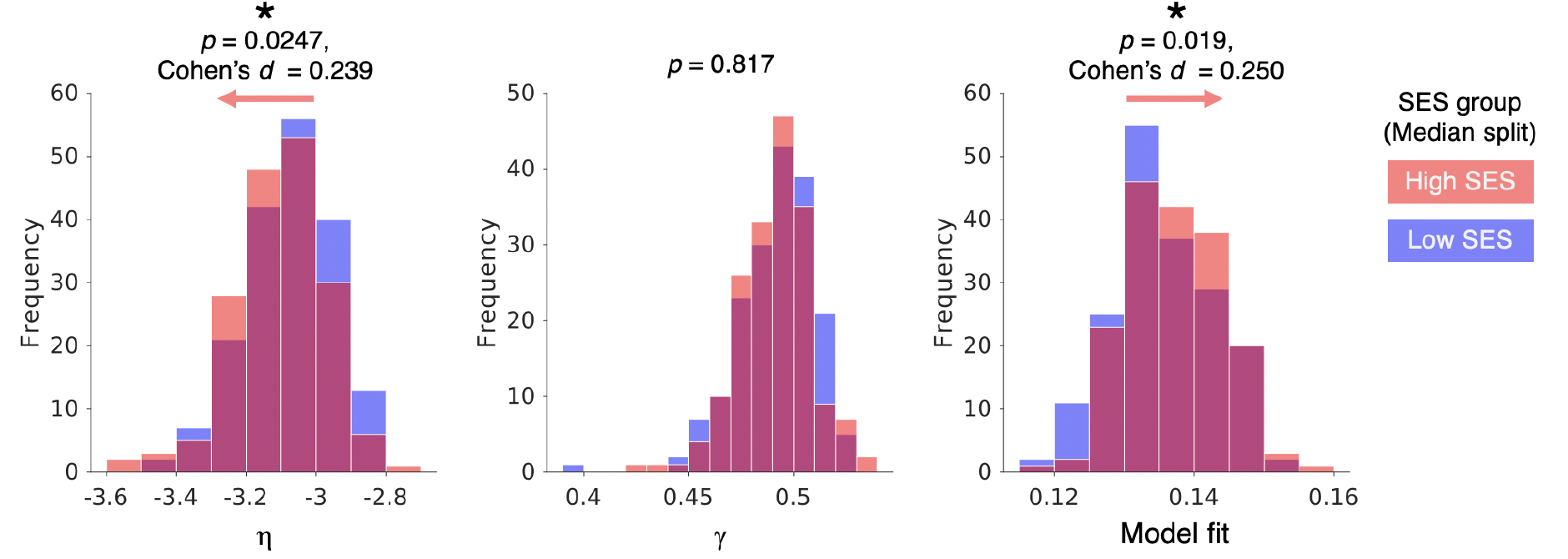
Wiring parameters and model fits vary with SES. In *n* = 357 children, we split groups into high and low SES groupings. We find that the wiring parameter *η* is greater magnitude negative in high SES children, suggesting more constrained connectivity (left). We find no effect in the *γ* parameter between groups (middle). Model fits are better in low SES networks reflected by being shifted to the left, due to the connectome being more randomly organized and therefore more easily simulated by the generative model (right).

In summary, heightened stochasticity within network development has a protective effect on the structure of the network, just as it protects against the impact of increased developmental noise (as shown in **Experiment 2**). However, this resilience is relative, rather than absolute.

### Experiment 4 – The effects of constraints on equifinality

We have so far established that weakening wiring constraints leads to greater multifinality in network outcomes and robustness to external perturbation, while highly constrained networks tend to have more invariant outcomes along with a greater vulnerability to change. Before turning to empirical data, we aimed to examine the relationship between stochasticity and a final core developmental principle: *equifinality*.

Equifinality refers to the attainment of a similar phenotype by way of a diverse set of pathways^34^, and is a known characteristic of child neural development^37^. Numerous contributing factors, such as similar environments and experiences, may contribute to similar phenotypes between two genetically different individuals.

We hypothesized that both the wiring constraints and the intrinsic stochasticity of the developing brain could modulate the resultant equifinality by setting the range of outcomes that any individual could exhibit. In other words, we predicted that brains generated using relatively similar wiring parameters would exhibit greater overlap in the range of potential phenotypic outcomes than those generated using more dissimilar parameters. However, we also expected that equifinality would depend on the intrinsic stochasticity of the models. Namely, we thought that simulations with weaker constraints would achieve lower equifinality with other parameter combinations than more constrained simulations, due to the broader range of potential outcomes of the simulation.

To test this prediction, we assessed the ability of a supervised machine learning model to successfully distinguish simulations run with differing wiring parameters based on their global topology. Specifically, we trained a support vector machine (SVM) to distinguish all possible pairwise comparisons of the global statistics of the simulations (see **Methods: Experiment 4**). Using 10-fold cross-validation, we computed the misclassification rate of the SVM. A higher rate would indicate that the SVM was performing closer to chance, and that the two sets of simulations were exhibiting equifinality. Lower misclassification rates would indicate that the SVM was successfully able to distinguish the two sets of simulations, and when this rate nears zero, the two sets could not be said to exhibit any equifinality.

As predicted, simulations run with closer generative modelling parameters tended to show higher equifinality (**Figure 5a**, *r* = -0.379, *p* < 0.0001). This reflects that a similar trade-off between wiring cost and value results in simulations that occupy an overlapping range of phenotypes (**Figure 1a**). Furthermore, simulations with higher multifinality in network topology also exhibited lower mean misclassification rate (**Figure 5b**, *r* = -0.787, *p* < 0.0001), indicating that developmental stochasticity was inversely related to equifinality. Thus, it appears that networks that grow less deterministically are more likely to end up in unique phenotypic outcomes that are easily distinguishable from more normatively developing networks.

### Prediction – Environmental uncertainty may favor weaker wiring constraints

Experiment 1 (*The effects of constraints on multifinality*) established that weaker wiring constraints lead to greater multifinality, whilst stronger wiring constraints narrow the range of phenotypic outcomes of the network. Experiment 2 (*The effects of timing on multifinality*) demonstrated that weaker wiring constraints are protective against the impact of heightened developmental noise, whereas stronger constraints allow noise to produce a time-dependent increase in multifinality. Experiment 3 (*The effects of constraints on robustness*) revealed that weaker constraints have a protective effect on the structure of the network, operationalized as resilience to relative change in communicability in response to targeted and random simulated attacks. However, this resilience is relative, rather than absolute. Finally, Experiment 4 (*The effects of constraints on equifinality*) shows that networks developing more stochastically more often exhibit strongly unique phenotypes, meaning there is a lesser likelihood of equifinal outcomes. Together, our findings reveal that stochasticity is a critical element of the successful simulation of brain networks, which may correspond to real-world developmental processes.

Our theoretical-computational framework gives rise to falsifiable hypotheses about child development. Namely, we propose that children growing up in unpredictable contexts (i.e., environments with fewer statistical regularities) have brain networks whose organization is best approximated with weaker wiring constraints (i.e., smaller magnitude *η* and/or *γ*). As per Experiment 1, this would correspond to more stochastic development. As per Experiments 2 and 3, such heightened stochasticity would make brain networks less sensitive to perturbation, which is likely a useful feature in uncertain environments. Finally, as Experiment 4 demonstrated, this up-regulation of stochasticity could lead to a greater likelihood of aberrant neural phenotypes as a by-product.

To test this prediction, we obtained data from *n* = 357 the Centre for Attention, Learning and Memory (CALM) (see **Methods** for more detail and **Supplementary Table 1** for demographics). To approximate the unpredictability of the early-life environment, we measured the Index of Multiple Deprivation (IMD) of each child, which captures their relative socioeconomic disadvantage. We then split subjects into high and low deprivation SES groups using the sample median IMD (**Supplementary Figure 4**). Next, we constructed structural connections of each child’s connectome using diffusion imaging and probabilistic tractography with anatomical constraints computed the best estimate of each child’s ground-truth wiring parameters (see **Methods**). In line with our prediction, we found that subjects with high SES show higher (negative) wiring parameter magnitude in *η* (**Figure 6, left**; *p* = 0.0247, Cohen’s *d* = 0.239). We found no detectable difference in the *γ* parameter (**Figure 6, middle;** *p* = 0.817). Finally, we found a corresponding better model fit in lower SES children (**Figure 6, right**; *p* = 0.019, Cohen’s *d* = 0.250), which accords with the fact that parameters are simulating more randomly organized networks—a comparatively easy target (see ^29^).

In summary, we tested the idea that—as stochasticity confers both robustness to perturbation and intrinsic variability in phenotypic outcomes—children within low-SES environments show an adaptive preference for heightened stochasticity within the topology of their macroscopic brain networks. Our findings, which show that the connection length constraint *η* is weakened in children from low-SES environments, appears to be consistent with this hypothesis.

## DISCUSSION

Some of the most fundamental elements of developmental theory — equifinality, multifinality and adaptability — are among the hardest to study empirically. In this work, through generative network modelling, we were able to investigate the role of stochasticity in the emergence of macroscopic brain networks. Importantly, this computational framework provided a means to systematically study these core developmental concepts. Through our simulations, we found that weaker wiring constraints leads to greater multifinality in brain network phenotypes, less sensitivity to temporary increases in developmental noise and greater relative robustness to simulated attacks, and greater likelihood of atypical phenotypes. By fitting our models to empirical data, we found that children from low-SES environments appeared to follow this developmental pattern: their connectomes were better approximated through a more stochastic generative model with weaker constraints on long-distance connections.

### The adaptive stochasticity hypothesis

Our work has highlighted, at a computational level, several core effects of developmental stochasticity. For example, it affords relative robustness to perturbation. That is, while it lowers the absolute capacity to support network communication, it confers resilience to change. Developmental stochasticity also inoculates networks against the impact of temporary increases in noise. Finally, higher stochasticity results in greater multifinality and more distinct network outcomes.

Are these features advantageous or disadvantageous? This likely depends on the environmental context of the developing system. In an enriched environment that is statistically predictable, stochasticity may be disadvantageous because it introduces greater risk of unfavorable outcomes that are unsuited to that narrow context. In contrast, networks developing within a statistically unpredictable environment benefit from what stochasticity can provide: flexibility to unexpected perturbation and robustness to change.

We therefore put forward the *adaptive stochasticity hypothesis*. This states that heightened stochasticity within the developing brain may serve as an adaptive mechanism in situations of environmental uncertainty. Our empirical finding that the brains of children from low socioeconomic backgrounds — which we use as an approximate measure of environmental predictability — are better simulated with more stochastic models offers preliminary and partial support for this hypothesis.

### Predictions

The adaptive stochasticity hypothesis gives rise to numerous testable predictions. If unpredictability increases developmental stochasticity in brain networks:

- *Connectomic variability*. Connectomic phenotypes should vary more amongst children who live in unpredictable environments.
- *Separable intrinsic and extrinsic influences*. Intrinsic (e.g., genomic) and extrinsic (e.g., environmental) factors should predict at least partially non-overlapping variance in wiring parameters.
- *Topological signature of randomness*. The up-regulation of developmental stochasticity should be manifest in network topology. Likely candidates for a measurable signature of stochasticity include lower segregation and greater integration^38^, consistent with higher entropy.
- *Elevated likelihood of non-normative outcomes*. Weaker wiring parameters, giving rise to heightened neurodiversity, should be associated with increased rates of neurodevelopmental conditions, captured by canonical diagnostic groups (e.g., schizophrenia^31^) or transdiagnostic dimensions^26^.
- *Adaptive cognition*. Weaker wiring parameters should be associated with adaptive outcomes on the cognitive level, including better performance on tasks that are relevant in harsh environments or under adverse testing conditions^39^. To our knowledge, no diffusion imaging cohort has yet collected such measures.
- *Inducible stochasticity*. Children raised in predictable environments later introduced to severe unpredictability should exhibit a shift in wiring parameters over time, depending on the chronicity of that unpredictability.

### Limitations

Our study has limitations in both methods and scope. First, our generative network modelling framework is a blunt approximation of stochasticity in the developing brain. Our models currently only approximate the topology of binary networks; new models that modulate connection strength over time may better capture developmental stochasticity. Secondly, this work does not consider the valence of the environment, which is entangled with but theoretically separate from unpredictability^38^. Moreover, we do not deal with meta-predictability, or the consistency of unpredictability over time^22^. Fine-grained measures of the environment and sophisticated modelling would be necessary to test how these may interact with developmental stochasticity. Finally, we have focused our analysis on macroscopic neural networks derived from diffusion tensor imaging, which is characterized by certain empirical limitations^40,41^ and cannot capture circuitry at lower scales.

### Conclusions

Stochasticity is an underappreciated contributor to developmental outcomes that may be particularly relevant in adverse environments. We present a modelling approach to parsing its contributions to the emergence of brain network organization. We believe this line of investigation could yield critical insights into inter-individual variability in general, and the influence of the early environment in particular.

## ACKNOWLEDGEMENTS

This publication is based on research supported by the Templeton World Charity Foundation, Inc. (funder DOI 501100011730) under the grant TWCF-2022-30510. Danyal Akarca and Duncan Astle are supported from the John S. McDonnell Foundation Opportunity Award and the Medical Research Council. Sofia Carozza is supported by the Cambridge Trust. The authors would like to thank Alexa Mousley for providing seed networks derived from the Developing Human Connectome Project (*dHCP*). For the purpose of open access, the author has applied a Creative Commons Attribution (CC BY) licence to any Author Accepted Manuscript version arising from this submission.

## CODE AND DATA AVAILABILITY

The empirical dataset supporting the current study have not been deposited in a public repository because of restrictions imposed by NHS ethical approval, but are available from the corresponding author on request. Requests for access can be made by research-based institutions for academic purposes. A response can be expected within 1 week.

Results were generated using code written in Matlab 2020b. Simulations were conducted on large compute clusters for parallelisation. All code will be made openly available upon publication.

## METHODS

### Probabilistic wiring equation

To simulate the formation of brain network connectivity, we employed generative network modelling^23,24^. This model is composed of a wiring probability equation:

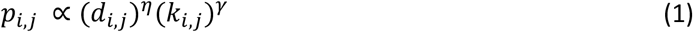

The *d*_*i,j*_ term represents the cost of a connection between two nodes *i* and *j*, approximated using the Euclidean distance between the two nodes. The *k*_*i,j*_ term represents how nodes *i* and *j* value each other and is set *a priori* using a topological relationship between the two (also denoted a “wiring rule”). Two wiring parameters, *η* and *γ*, respectively parameterize the costs and value terms, thereby calibrating their relative influence. *p*_*i,j*_reflects the probability of forming a fixed binary connection between nodes *i* and *j*. This is proportional to the parametrized multiplication of costs and values.

Previous research has shown that generative models implementing a value term (i.e., wiring rule) that prefers connections between regions with overlapping neighbors, termed *homophily*, can reliably produce networks with statistics that mirror empirical observations^25,26,28,29,42^. As such, we utilized the matching wiring rule in our analyses – the main homophily wiring rule used in prior work.

Mathematically, matching is defined as the normalized overlap in connectivity between two nodes^24,25^. Specifically, if Γ represents the set of node *i*’s neighbors, then the matching index is equal to:

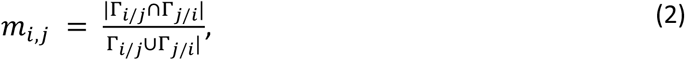

where Γ_*i*/*j*_is Γ but with *j* excluded from the set. If *i* and *j* have perfect overlap in their neighborhoods, then *m*_*i,j*_= 1. If the neighbourhoods contain no common elements, then *m*_*i,j*_= 0. This is because ∩ (the numerator) refers to the *intersection* of the neighbors (i.e., neighbors in common) while ∪ (the denominator) refers to the *union* of the neighbors (i.e., neighbors in total).

All simulations were generated within a physical space defined by the commonly-used AAL116 parcellation scheme^43^. The coordinates of node centroids within the AAL116 atlas were used to determine the Euclidean distance between every node combination, which was used to approximate the cost of connections.

A termination criterion must be defined for when the formation of the networks ends. As we do not mirror empirical data for the main part of the study, we set 400 connections as the stop criterion to attain a final density of 3%. This was done so that the core trade-offs could easily be examined within computational limits. Later in the study, when we compare directly to empirical data, we use the number of edges in the empirical *CALM* dataset networks (see **Methods: Empirical prediction and application**) as the stop condition. This gives at an average sample 10% density (mean 668.5 edges; SD 43.6 edges), which aligns with prior work done on the sample^26^ and elsewhere^25^.

### Neonatal seed network generation

The approach to initializing generative network models (i.e., the seed) that best mimics developmental trajectories is unknown. Three main approaches have been taken in the literature: (1) The selection of edges that are highly consistent across the sample of empirical connectomes^25,26,29^; (2) No edges at all, thereby initializing the model with a first calculated *p*_*i,j*_matrix that is equivalent to *d*_*i,j*_matrix^24,28,30,44^ and; (3) A theory-driven set of edges (e.g., medio-posterior nodes^23^).

The choice of initial conditions is of great biological relevance. The generative model is thought to reflect activity-dependent interactions between neural assemblies by forming connections between self-similar regions. However, it is widely known that the preliminary scaffold of brain connectivity arises through processes that are largely activity-independent^45^. Furthermore, by virtue of wiring early on in development, these regions will have a greater time-availability for future wiring therefore are more likely to become hubs later in development (the old-get-richer effect^46^). We sought to account for the early activity-independent scaffold using a neonatal seed network.

Toward this end, we reconstructed a core rich club network from data collected from the Developing Human Connectome Project (*dHCP*). We then used this core neonatal rich club as the initial connectivity matrix for our subsequent simulations. The *dHCP* sample contains *n* = 630 neonates (mean post-conceptual age = 39.46 weeks, SD = 3.58 weeks, *n* = 297 female, *n* = 343 male) connectomes rendered within the AAL90 parcellation (^47^; see **Methods: Neuroimaging data and preprocessing**). A rich club topology describes how high-degree nodes tend to be more densely interconnected (in topological binary networks) than would be expected by chance.

To identify highly conserved invariable edges across the *n* = 685 neonatal connectomes, we considered only those edges which were shared in 70% of the sample (*n* = 480) (**Supplementary Figure 1a, b**). Across the sample, these edges had above-average streamline connectivity (**Supplementary Figure 1c**).

To assess the inter-connectivity between hub regions within a binary brain connectivity network, we used the topological rich-club coefficient *ϕ*(*k*). This quantifies the density of the subgraph comprising nodes with a degree higher than the hub-defining threshold *k*:

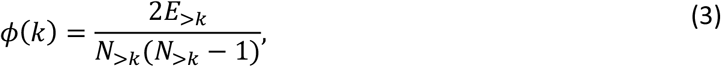

where *N*>*k* is the number of nodes with degree>*k*, and *E*>*k* is the number of edges between nodes with degree >*k*.

Because higher degree nodes make more connections, it is important to determine whether this value is higher than that expected by change. We therefore compared the rich club coefficient of the empirical neonatal consensus network to the mean value across a 1000 randomized null networks, generated by rewiring the edges of the empirical network while retaining the same degree sequence, using the *randmio_und* function from the Brain Connectivity Toolbox^48^, rewiring each edge 50 times per null network. This approach has precedent in the literature (e.g., ^44^). We thus computed a normalized rich-club coefficient, taking the ratio between the rich-club coefficient in the empirical network and the mean rich-club coefficient in this set of corresponding randomized networks:

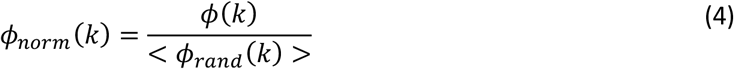

Values of >1 indicate rich-club organization, where high-degree nodes are more densely interconnected. To characterize the statistical significance of the result, we computed a *p*-value directly from the empirical null distribution of the 1000 randomized networks, *ϕ*_*rand*_(*k*), as a one-sided permutation test. Specifically, the *p*-value is the ranking position of the empirical rich-club coefficient within the null distribution of rich-club coefficients from the randomization procedure. For example, a *p*_*perm*_ of 0.01 would be equivalent to being in the top 1% of 1000 null models, which is position 10.

In **Supplementary Figure 2a** we plot the rich-club coefficient, normalized rich club coefficient and p_perm_ values across increasing degree, or *k* levels. This highlights a rich-club topology that is statistically significant (*p* = 0.018) is found beyond level 22. That is, there is a sub-graph of nodes with degree>22 in the neonatal consensus network that are connected to each other more than would be expected by chance. This rich club network contains 5 nodes: left and right lingual cortex, left precuneus, left occipital mid and right fusiform cortex. All nodes fully connected to each other (*n* = 10 edges) (**Supplementary Figure 2b, left**). This seed network generally occupies medio-posterior positions which is thought to be the earliest location of white and grey matter development^23^.

To scale these results to adult size, we then placed this equivalent rich club within the adult AAL116 equivalent atlas (**Supplementary Figure 2b, right**). This network was then used as the seed for all simulations. Note that, to prevent data-leakage, we did not assess model fits with respect to any of the *dHCP* data. Instead, we only tested models on a completely independent dataset (see below).

As the seed network consists of 10 edges, and simulations were computed for 400 edges (density = 0.075%), the seed constitutes the first 2.5% and 1.05% of simulated networks in the main analyses versus empirical analyses respectively. As seen in **Supplementary Figure 2a**, a lower-level threshold of degree >17 would also generate a significant rich club network, but this constituted 133 edges and therefore was deemed too large.

### Parameter space and repeated simulations

We ran the generative models from this seed network at 625 different parameter combinations of *η* and *γ*. The parameter combinations were selected evenly across a parameter space of 0 ≤ *η* ≤ 4 and 0 ≤ *γ* ≤ 1 to provide a 25×25 grid space. This parameter space was chosen as this range best recapitulates the statistics of empirical brain networks using the matching rule^25,26^. At each combination, we ran the simulation 625 times. This led to a total of 390,625 (625 parameter combinations x 625 repetitions) simulations for each protocol.

### Experiment 1 – Network outcome dissimilarity

For each parameter combination, we computed two measures of dissimilarity between every combination of the produced 625 networks (**Figure 1b**). The first dissimilarity measure was a measure of topological dissimilarity. For each network, we calculated five simple global topological measures for each network, using the Brain Connectivity Toolbox^48^: (1) Global clustering; (2) Mean betweenness centrality; (3) Total edge length; (4) Global efficiency and (5) Modularity. These were used because they cover a range of topological features common to brain networks. From this topological matrix, we then simply calculated the Euclidean distance between each network in this 5-dimensional space (625 × 625 dissimilarity matrix). As measure of total topological dissimilarity, we simply computed a summation of this Euclidean distance matrix (**Figure 1c**). The second dissimilarity measure was a measure of embedding dissimilarity, which calculated the percentage of non-overlapping connections (i.e., the dis-consistency) between all network combinations (**Figure 1d**). For both measures, large numbers correspond to there being more heterogeneity among network outcomes, and *vice versa*.

### Experiment 2 – Timing of noise analysis

To assess the effects of heightened stochasticity in network development on variability in outcomes, we re-ran each of our models whilst forming a subset of the connections at random. This was achieved by temporarily setting the probability matrix to zero, erasing any contribution made by the cost and value of connections and allowing network formation to proceed completely randomly. We injected this noise at one of three stages of development: early, middle, and late. This corresponds to 5-10%, 47.5%-52.5%, and 90-95% of network development. We then computed the global topological dissimilarity (see **Methods: Experiment 1**) of the resulting networks. This enabled us to identify and quantify the contribution of injecting noise at the three points in development to the stochasticity of network outcomes.

### Experiment 3 – Robustness analysis

Network robustness refers to the ability for networks to be resistant to external perturbation. It is evaluated by computing some benchmark measure of network quality before and after perturbation. As previous studies suggest that communication models accurately capture propagation dynamics in empirical brain networks^49,50^, we selected binary network communicability^51^ as the benchmark measure:

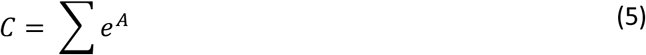

Here, *A* is the symmetrical binary matrix that has been simulated and *C* is the network communicability within that matrix.

We used two regimes to assess robustness of the networks: (i) targeted and (ii) random attacks. In the targeted attack regime, nodes within a network are first ranked according to some measure (in our case, the degree of the nodes). Then, nodes are incrementally removed (i.e., attacked) by tuning all their connectivity weights to zero. In the random attack protocol, nodes chosen at random are attacked in the same way. After each node’s connectivity is removed, network communicability was recalculated.

Robustness—or resistance to change—was subsequently measured as the coefficient of a univariate linear model fit to the trajectory of the natural logarithm of the communicability after 25% of total nodes (29 nodes). This indicates the extent to which a network is robust to change in its capacity to support dynamics; the larger in magnitude the negative coefficient, the less robust the network, and *vice versa*.

### Experiment 4 – Equifinality analysis

To test which factors may contribute to equifinality, we assessed the ability of a supervised machine learning model to successfully distinguish simulations run with differing wiring parameters. For each of the previously-run 625 simulations, we trained a support vector machine (SVM) to distinguish its global statistics from each of the 624 other simulations. Then, we used 10-fold cross-validation to determine the misclassification rate (i.e., how many of the 625 runs were attributed to the wrong parameter combination). In order to determine the contribution of wiring parameters to equifinality, we calculated the Pearson correlations between the misclassification rate and the Euclidean distance between the two parameter combinations. To determine the contribution of the intrinsic stochasticity to equifinality, we calculated the Pearson correlation between the mean misclassification rate (across the 624 comparisons) and the topological stochasticity of the simulation (see **Methods: Experiment 1**).

### Empirical prediction and application

To test our theoretical framework, we made empirical predictions about the relationship between socioeconomic status and brain wiring parameters, which we then tested in a large sample of children.

#### The CALM cohort

The data were collected at the Centre for Attention, Learning and Memory (CALM), a research clinic at the MRC Cognition and Brain Sciences Unit, University of Cambridge. The study protocol was approved by, and data collection proceeded under the permission of, the local NHS Research Ethics Committee (reference: 13/EE/0157). The sample is designed to be reflective of children at heightened risk of a range of neurodevelopmental difficulties^52^ and has sufficient variability within it to establish wiring parameter differences^26^. Practitioners working in specialist educational or clinical services in the East of England (UK) were asked to refer children with ongoing problems of “language”, “attention”, “memory”, or “learning/poor school progress”, regardless of the presence or absence of a formal diagnosis. Exclusion criteria included uncorrected problems in vision or hearing, having English as a second language, or having received a causative genetic diagnosis. In addition to 800 children, 200 children recruited from the same schools and neighbourhoods who present no such difficulties. A range of measures were collected, including genetic, cognitive and behavioural, and neural data (see ^52^ for the full assessment protocol). Of the *n* = 1000 total children, *n* = 425 completed some neuroimaging portion of the study, of whom *n* = 386 had diffusion imaging data (see below for pre-processing details). A strict movement threshold (see below) led to a final sample of n = 357. Of these children, *n* = 283 were from the pool of 800 children meeting the above *CALM* criteria, and *n* = 73 were from the control group.

#### Neuroimaging data and preprocessing

MRI data were acquired on a Siemens 3 T Prisma-fit system (Siemens Healthcare) using a 32-channel quadrature head coil. T1-weighted volume scans were acquired using a whole brain coverage 3D Magnetization Prepared Rapid Acquisition Gradient Echo sequence acquired using 1 mm isometric image resolution. Echo time was 2.98 ms, and repetition time was 2250 ms. Diffusion scans were acquired using echo-planar diffusion-weighted images with an isotropic set of 68 noncollinear directions, using a weighting factor of b = 1000 s mm^−2^, interleaved with 4 T2-weighted (b = 0) volume. Whole brain coverage was obtained with 60 contiguous axial slices and isometric image resolution of 2 mm. Echo time was 90 ms and repetition time was 8500 ms. We enforced a strict movement threshold of 1mm (estimated through FSL eddy during the diffusion sequence), which led to 29 scans being removed, leaving a final sample of 357 children.

Images were preprocessed using FSL eddy to correct for motion, eddy currents, and field inhomogeneities. Nonlocal means de-noising using DiPy v0.11 was performed to increase signal-to-noise ratio. Finally, single-shell constrained spherical deconvolution (CSD) was used to estimate the fibre orientation distribution^53^ with a brain mask derived from the T1-weighted image. Whole-brain tractography was then performed using iFOD2 probabilistic tracking^54^, to generate 10^7^ streamlines with a maximum length of 250mm and minimum length of 30mm. Weights for each streamline were calculated using SIFT2^55^. Streamlines were then mapped to the Schaefer 100 atlas, a gradient-weighted Markov Random Field parcellation of the cortex^56^, to generate a connectivity matrix.

We thresholded this connectivity matrix at an absolute threshold of 1530 streamlines to generate binary connectomes for our analyses to achieve a sample average 10% of density (see **Methods: Probabilistic wiring equation** for details on the number of connections present across the sample). We then simulated the development of each structural connectome using the generative network modelling procedure as outlined above, and determined the parameters that best replicated their organization through the Fast Landscape Generation (FLaG) approach described below.

#### Fitting generative model parameters using Fast Landscape Generation (FLaG)

A significant shortcoming of generative modelling in large cohort studies is that parameter estimation is computationally burdensome. Here, we use a fast, reliable, and accurate parameter estimation method for connectome generative models called Fast Landscape Generation (FLaG; ^33^). This method computes multiple landscapes and fits subjects to individual simulations simultaneously. At our sample size of *n* = 357, this allows for efficient but clear examination of differences. As recommended, we computed *k* = 50 landscapes to compute the average parameter combination across landscapes per subject.

#### Analysis of socioeconomic status

To measure socioeconomic status, we used the Index of Multiple Deprivation (IMD). This index captures the relative deprivation of an individual’s circumstances by surveying income, employment, education, health, crime, barriers to housing and services, and the quality of the living environment. See **Supplementary Figure 3** for the distribution of IMD values in the CALM sample.

