## Supplement for "The adaptive stochasticity hypothesis: modelling equifinality, multifinality and adaptation to adversity"

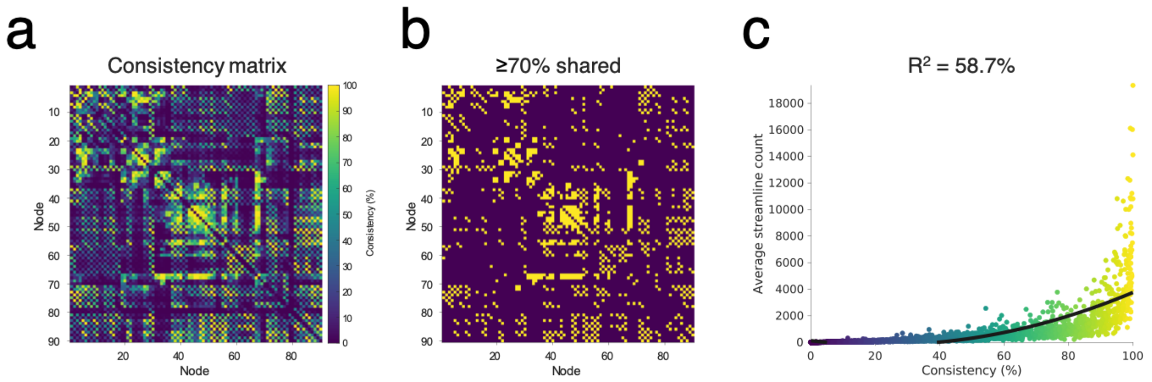


**Supplementary Figure 1. Thresholding *dHCP* neonatal connectomes through consistency-based thresholding of the *n* = 685 *dHCP* neonatal connectomes.** (a) The consistency matrix of how common an edge is across the sample. (b) The connectivity matrix, filtered to include only those edges which exist in ≥ 70% (*n* ≥ 480 of the sample). (c) The consistency of the edge across the sample is positively related to the average streamline count of that edge. A quadratic model fit to the data gives $R^2$ = 58.7%.

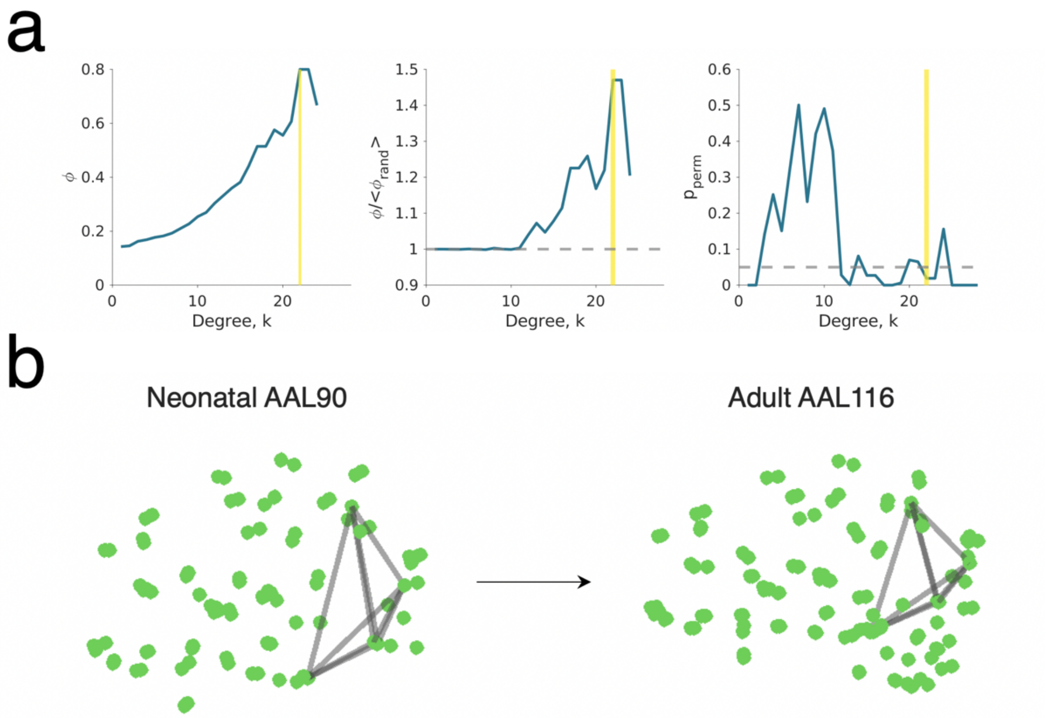


**Supplementary Figure 2. Neonatal rich club topology as the generative model seed.** (a) The rich-club coefficient chart for increasing degree, k thresholds. The selected threshold of $k$ = 22 is highlighted, showing the highest $\phi$ = 0.8 (left). The normalized rich-club coefficient chart for increasing degree, $k$ thresholds. Normalization was performed using the mean of 1000 randomized degree-preserving null models. The selected threshold of $k$ = 22 is highlighted showing the highest $\phi/<\phi_{rand}>$ = 1.5 (middle). The permuted $p$-value ($p_{perm}$) chart for increasing degree, $k$ thresholds. The selected threshold of $k$ = 22 is highlighted showing significance; $p_{perm}$ = 0.016 (right). (b) A graphical representation of the neonatal rich club in AAL90 MNI space, used as the seed network for generative modelling.

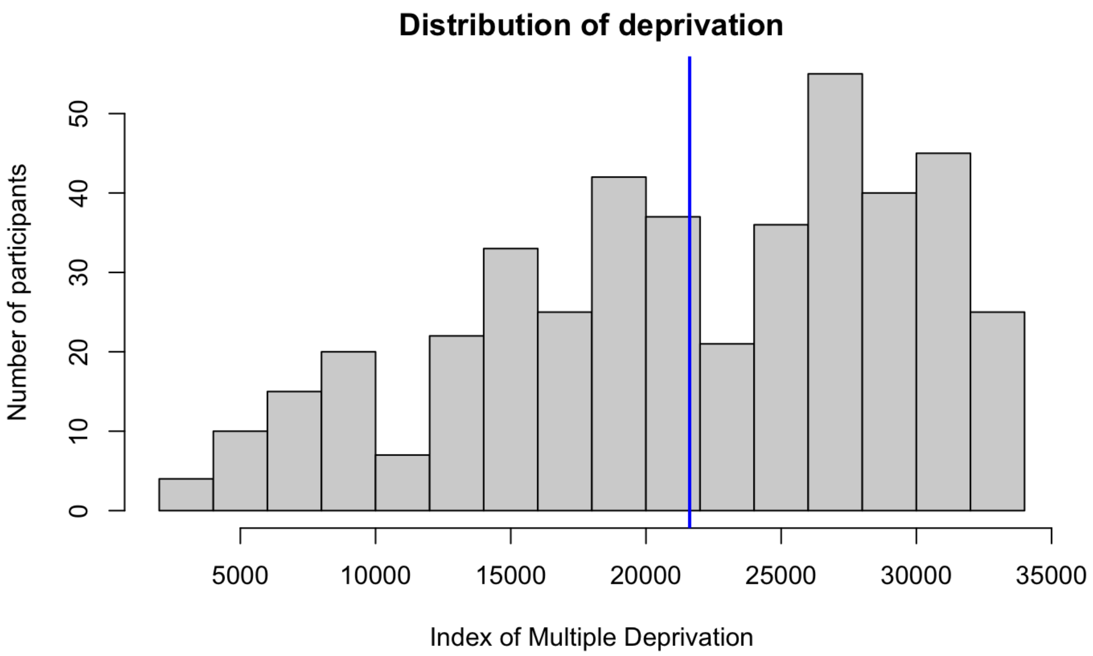


**Supplementary Figure 3.** Distribution of values of the index of multiple deprivation (IMD), a measure of the relative deprivation of the participant's area. The IMD accounts for income, employment, education, health, crime, barriers to housing and services, and the quality of the living environment. The blue line indicates the mean of the sample.

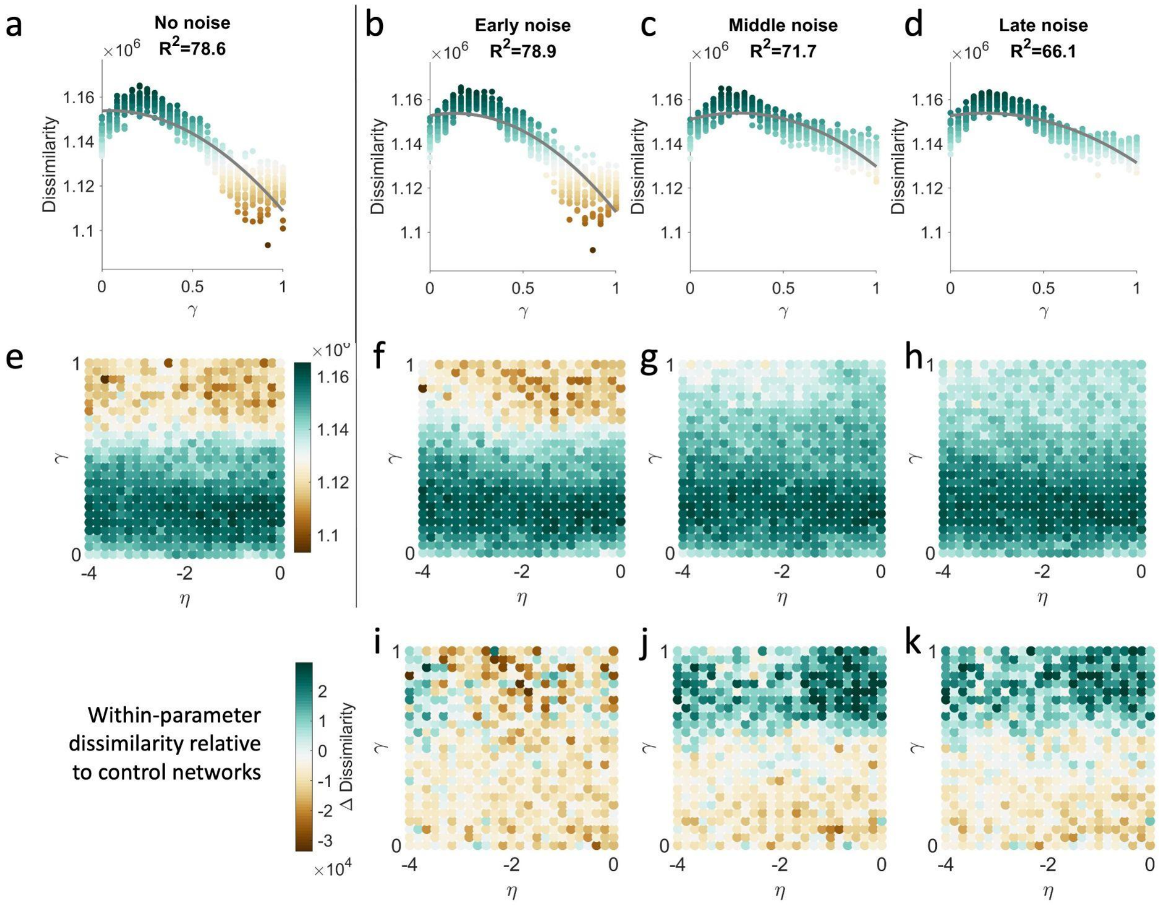


**Supplementary Figure 4. Stochasticity later in network development results in greater variability in network topology.** Compared to simulations run without noise (a) simulations with noise injected exhibit greater topological dissimilarity. Compared to (b) early noise, (c) middle and (d) late noise have a greater impact. When considering a heatmap of dissimilarity across the parameter space, the greatest increase in dissimilarity is localized at higher values of $\gamma$ (e-k).

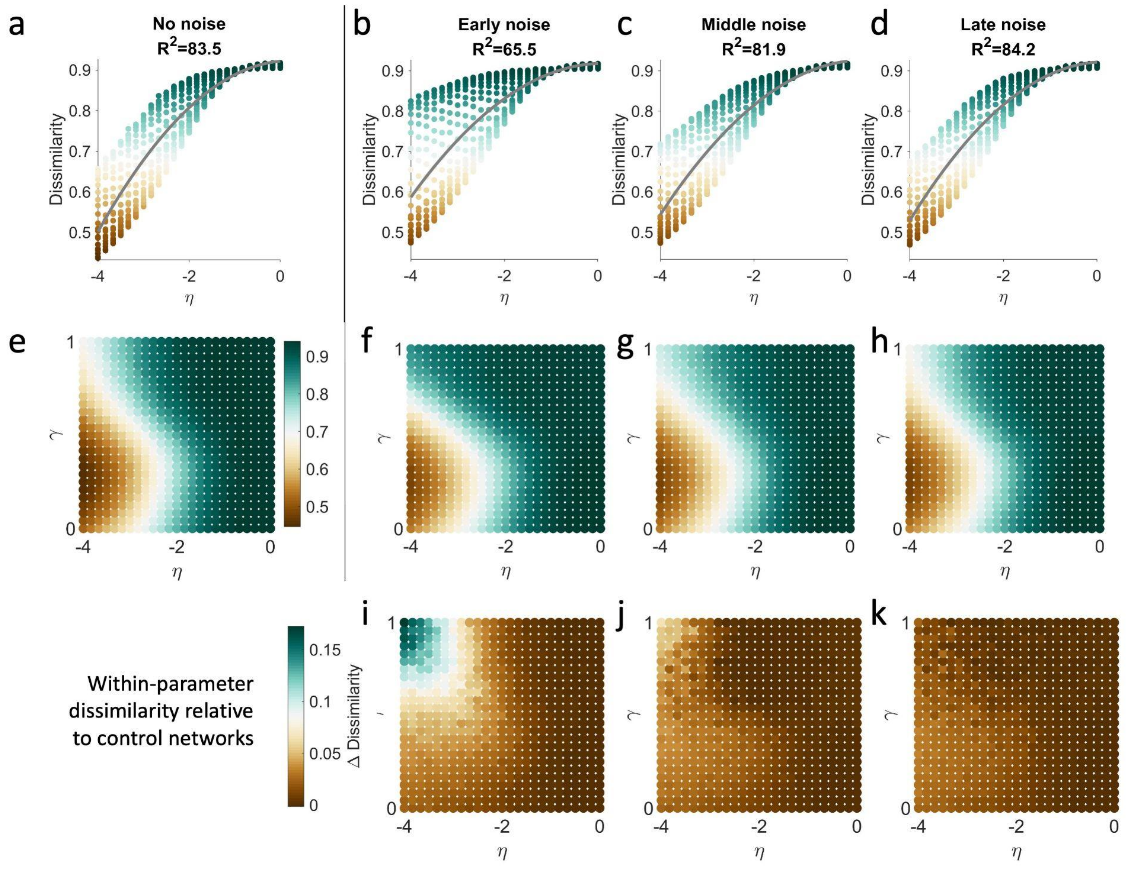


**Supplementary Figure 5. Stochasticity later in network development results in greater variability in the consistency of connection.** Compared to simulations run without noise (a) simulations with noise injected exhibit greater dissimilarity in embedding. Compared to (b) early noise, (c) middle and (d) late noise have a lesser impact. When considering a heatmap of dissimilarity across the parameter space, the greatest increase in dissimilarity is localized at lower values of $\eta$ (e-k).

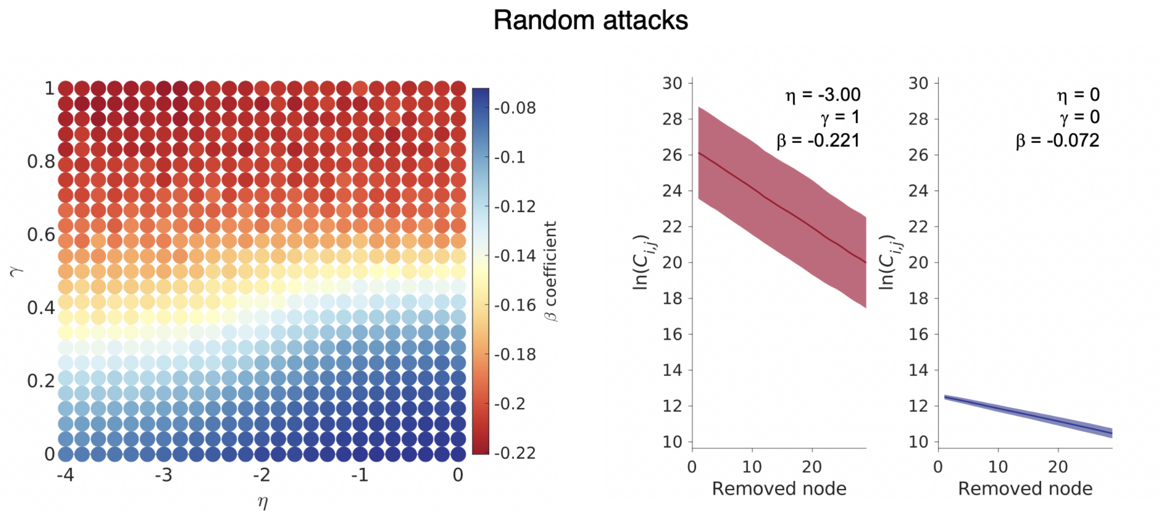


**Supplementary Figure 6. Robustness trade-offs in network connectivity following random attacks.** The $\beta$ coefficient computed from a random attack regime is provided, showing that higher parameter (top left of the left landscape) networks have less robustness as the gradient of change is greater and vice versa (bottom right of the left landscape). Example network regimes are highlighted to the right, showing outcomes with the least (left) and most (right) robustness to change.

| | Pre-Imputation ($n$ = 437) | Post-Imputation ($n$ = 437) |
|---|---|---|
| | **Age (months)** | |
| min | 62 | 62 |
| max | 223 | 223 |
| mean(sd) | 118.73 ± 27.81 | 118.73 ± 27.81 |
| | **Sex** | |
| Male | 285 (65) | 285 (65) |
| Female | 152 (35) | 152 (35) |
| | **Index of Multiple Deprivation** | |
| min | 2208 | 2208 |
| max | 32840 | 32840 |
| mean(sd) | 21,635.58 ± 7,795.45 | 21,615.05 ± 7,711.62 |

**Supplementary Table 1.** Descriptive statistics of the demographics of the CALM cohort.
